# Haptic perception is contingent to hemispaces, not to hands

**DOI:** 10.1101/2024.06.04.597475

**Authors:** Jess Hartcher-O’Brien, Alexander V. Terekhov, Vincent Hayward

**Affiliations:** Meta Reality Labs, Redmond, Washington, USA; Redwood Center For Theoretical Neuroscience, University of California, Berkeley, California, USA; Sorbonne Université, Institut des Systèmes Intelligents et de Robotique, Paris, France; Institute of Philosophy, School of Advanced Study, University of London, London, UK

## Abstract

It is known that human haptic perception is lateralised, for example, object shape is felt differently according to the hand used to explore objects. Here we show that it is not the hand but the hemispace in which the exploring hand is located that determines differences in perception. This finding implies that our lateralised somatosensory processing depends on hand localisation in space rather than on the hand itself.

## Introduction

We employed a curvature judgement task to test haptic shape perception invariance across hemispaces and across hands. A solid shape is entirely described by its curvature, which is invariant under rigid transformations [1]. It follows that if curvature is not felt in an invariant manner across hemispaces or across hands, then shape would not be either. Object curvature is accurately gauged by humans through the variation of first-order geometric information acquired by the contact of fingers against continuous surfaces [2–4].

In a “matching-side” experiment, participants explored a standard curved surface located in one of the two hemispaces with the hand corresponding to that same hemispace and explored a comparison curved surface located in the contralateral hemispace with the hand corresponding to the contra-lateral hemispace, see Fig. 1a. In a “same-hand” experiment, participants explored a standard curved surface with their dominant hand and explored a comparison curved surface located in the contra-lateral hemispace with the same hand, see Fig. 1b. The participants” dominant hand therefore transferred from one hemispace to the other. The experiments gave rise to four conditions since the standard curved surface could be located either in the right or in the left hemispaces (see Methods).

**Figure 1:**
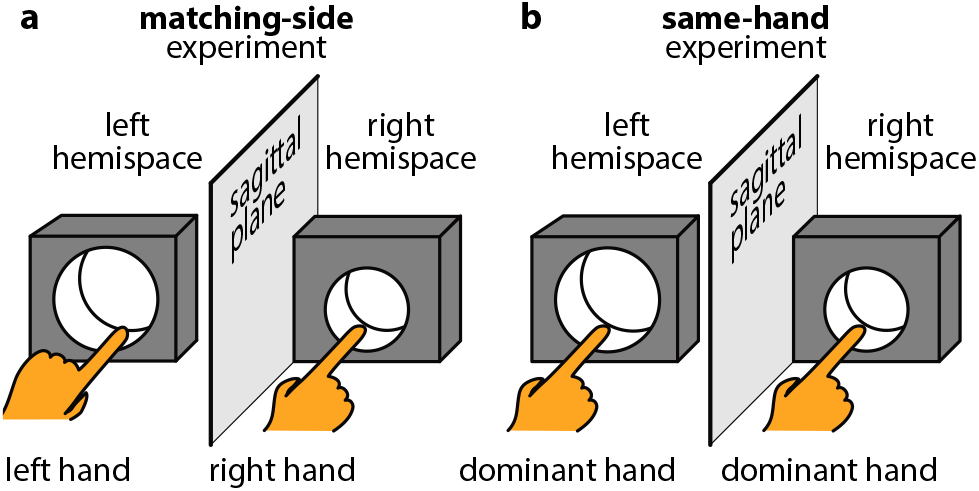
“Matching-side” and “same-hand” experiments. The standard surface had a curvature of 36 m^−1^ and could be located in the right or the left hemispace. **a**, “matching-side” experiment (*n* = 16). **b**, “same-hand” experiment (*n* = 6).

We found that the human haptic shape of solid objects depends — not on the hand exploring the object — but on the region of external space in which the hand explored the object. Similar results were obtained with object stiffness and size judgments.

## Results

Point of Subjective Equivalence (PSEs) expressed the curvatures that were perceived to be the same as the standard of 36 m^−1^ (Methods). These averaged values across participants, representing the bias in perceived curvature, are reported in Fig. 2a,b. The experiments replicated previous results for curvature discrimination thresholds of concave shapes (e.g. [4]). There were no significant differences between the four conditions, “matching-side” experiment, *t*(1) = 0.63, *p* = 0.53; “same-hand” experiment, *t*(1) = 0.01, *p* = 0.91, aside from the fact that using a right hand, frequently the dominant hand, predictably led to lower discrimination thresholds when it was in the right hemispace.

**Figure 2:**
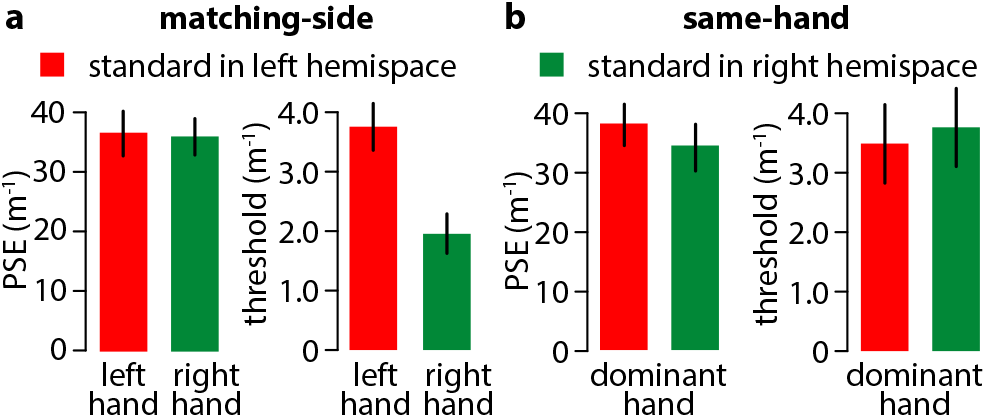
Results pooled across participants. **a**, mean PSEs and discrimination thresholds in the “matching-side” experiment. **b**, mean PSEs and discrimination thresholds in the “same-hand” experiment. Error bars show standard deviations.

In the “matching-hand” experiment we observed a surprising negative correlation on an individual basis between the biases in perceived curvature and lateralisation, see Fig. 3a, *R* = −0.88, *p* = 0.01. The participants who felt objects to have a greater, respectively smaller, curvature than the standard when using their left hand experienced the same object to have significantly smaller, respectively greater, curvature when exploring with their right hand. This negative correlation was also strong in the “same-hand” experiment where a given hand, right or left, was relocated in different hemispaces, Fig. 3a, *R* = −0.95, *p <* 0.01.

**Figure 3:**
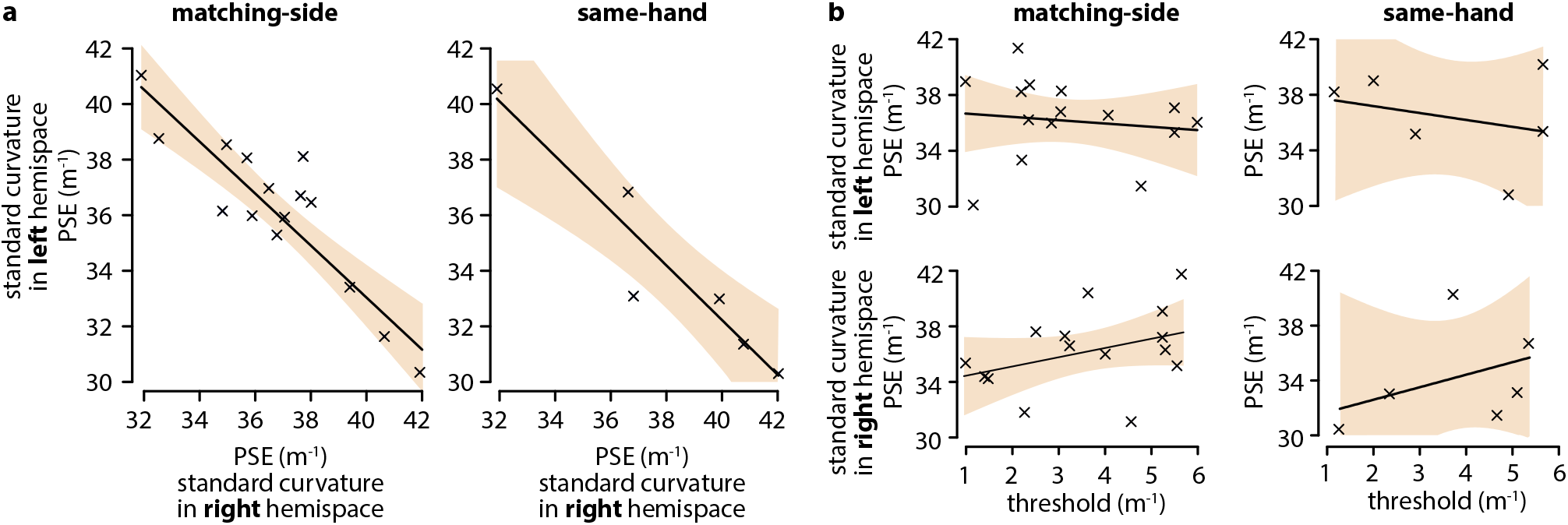
Individual results. **a**, right hemispace PSE vs left hemispace PSE estimates of perceived curvature in the two experiments. **b**, PSE vs thresholds estimates of perceived curvature in all four conditions. The light brown areas represent the 95 % confidence intervals of the fit of a first-order model.

In the “matching-side” experiment, a first-order regression across participants indicated that the bias in perceived curvature was linked to locating the left hand in the left hemispace and to locating the right hand in the right hemispace, *R*^2^ = 0.79, *F* (1, 15) = 52.87, *p <* 0.001. The same was true for the “same-hand” experiment, *R*^2^ = 0.89, *F* (1, 5) = 35.65, *p <* 0.01. The discrimination thresholds, in line with those previously reported in the literature [5], could not account for the biases in perceived curvature, see Fig. 3b, *R*^2^ = 0.1, *F* (1, 16) = 0.47, *p >* 0.05. Locating a hand in a given hemispace predicted the bias but the hand did not. On an individual basis, the data showed that the biases in perceived curvature arose from a hemispace-dependent coding of touch, rather than from a response bias.

Other object properties, namely stiffness and size, were perceived with a bias similarly correlated to the hemispace in which the exploring hand was located, see Fig. 4. This bias was also consistent across experiments, see Fig. 5. The data also showed that the biases in perceived curvature arose from a hemispace-dependent coding of touch, rather than from a response bias.

**Figure 4:**
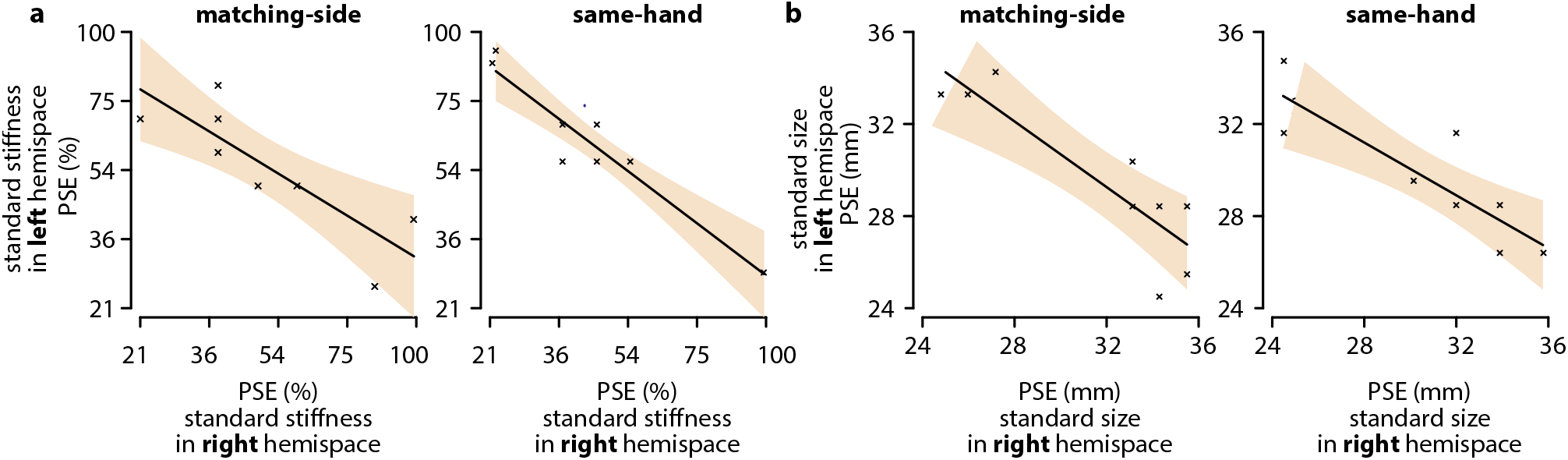
Individual results. **a**, condition where participants judged the stiffness of an object. Stiffness was gauged bi-digitally by squeezing a calibrated spring through a lever mechanism providing an adjustable mechanical advantage (Methods). The scale was quadratic and referred to a standard set at 54% of the calibrated spring stiffness (Methods). **b**, condition where participants judged the size of objects by palpating them bi-digitally (Methods). The standard size was 30 mm. The light brown areas represent the 95 % confidence intervals of the fit of a first-order model.

**Figure 5:**
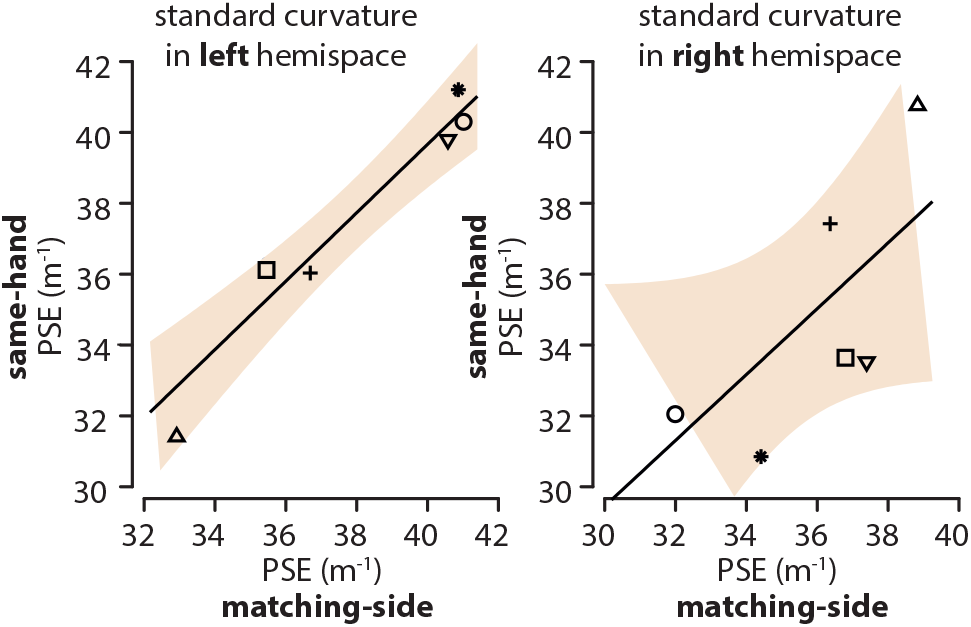
Individual participant’s PSEs across experiments. The light brown areas represent the 95 % confidence intervals of the fit of a first-order model.

## Discussion

The perceptual differences across space were not systematic across participants. For some participants, an object located in the ipsilateral hemispace was perceived to be more curved, more stiff, or larger than the same object located in the contralateral hemispace. For others, this pattern was reversed.

### Differences with previous research

Previous research on haptic shape perception was carried out with participants discriminating between curved profiles unimanually. Participants typically scanned two surfaces consecutively with the same hand (e.g., [4]). Engaging bimanually with objects provides redundant information to the brain about an object’s properties. Previous work on bimanual haptic shape perception suggests that haptic shape inputs across the hands are not integrated [6]. Moreover, bimanual interaction with a single object appears to decrease performance [5], highlighting a difference in processing between the visual and somatosensory systems.

### Examples of lateralised haptic perception

There are many examples of lateralised somatosensory perceptual effects. The “rubber hand illusion” does not extend to hands crossing the sagittal plane [7], and haptic line bisection is also affected by the position of the line in hemispace, but is independent of the hand that bisects the line [8]. Illusions of cutaneous apparent motion do not cross the body midline [9]. The perceptual asymmetries related to the location of objects in lateralised space may be due to the fact that communication across hemispheres is partial [10, 11], representing early lateralisation during development [12].

### Lateralisation in vision and touch

Perceptual lateralisation was not documented in vision where for most participants, changes in the proximal stimulation are invariant under a change of hemifield [13]. One reason for this difference may be that in human vision the stereopsis pathways share information about the same central visual field regions independently of the hemispace in which they are located, whereas in uni-manual haptic interactions, the hands can cross hemispaces but objects tend to remain fixed within their hemispaces. In vision, shape constancy is thought to be achieved by converting retinal signals in external coordinates, allowing the observer to perceive space independently of the movements of the eyes [14, 15].

Maintaining perceptual constancy in touch can be thought to be a harder problem to solve than in vision and may receive more varied solutions across individuals. Behavioural evidence further indicate that locating objects in different hemispaces can influence perceptual judgements [16–21]. There is compelling evidence for both inter-hemispheric and lateralised somatosensory processing [22, 23].

Our findings are consistent with a lateralisation model of somatosensory processing where the perceived form of an object is independent of the hand exploring the object but is linked to the region of external space that the hand and the object occupy [24].

## Methods

The curved stimuli were cylindrically holed-out blocks machined out of acetal plastic. The inner surfaces of the holes were finely sanded to an average roughness, *R*_a_, of about 10 μm so that they were smooth but not mirror finish, regularising the frictional properties of the finger-surface interface. The curvature of the standard stimulus was set at 36 m^−1^ and the comparison stimuli had curvatures ranging from 46 m^−1^ to 26 m^−1^ in steps of 2 m^−1^. The heights of the low points of the holes were the same for all stimuli, Fig. 6a, to control for a possible curvature cue [25]. Participants were instructed to keep the fingertip level, Fig. 6b. To gauge object stiffness, participants palpated bi-digitally a spring-loaded lever mechanism where a spring of stiffness *k* could be located at a distance *h* from the fulcrum. If *l* was the distance from the finger to the fulcrum, then the stiffness experienced by the participant was *kd*^2^*/l*^2^. Seven comparison stiffnesses were presented to the participants with *d*^2^*/l*^2^ ranging from 21% to 100% with the standard set at 51% of the calibrated spring stiffness *k*. To gauge object size seven wooden stick were cut at lengths from 24 mm to 36 mm by steps of 2 mm with the standard size set at 30 mm.

**Figure 6:**
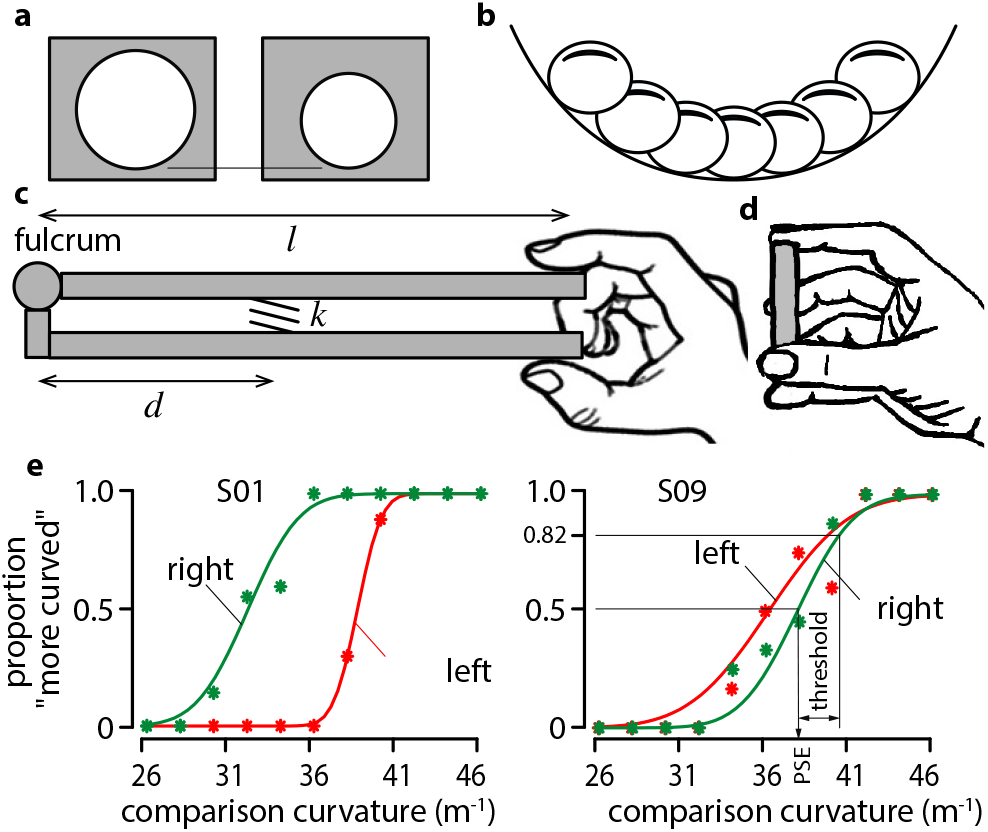
PSEs and thresholds.

Sixteen naive participants were recruited, aged between 19 and 35 years, in the “matching-hand” experiment and six participants were recruited in the “matching-side” experiment in the same age bracket. Participants where the same per condition expect one. All reported normal somatosensory sensations. They gave their informed consent prior to participation in the experiment. The testing procedures were approved by the “Comité de protection des personnes Ile-de-France II” permit 2011-06-16 (IRB registration 1072). Participants were blindfolded and sat in front of a table at elbow height. Two stimulus clamps were positioned 10 cm in depth from the table edge and 60 cm apart. The midline of the participants body was aligned with the workspace. In a two-interval, forced-choice procedure, participants used bare-finger contact to sweep or palpate six times each stimuli consecutively. They always returned their hand to the table between two scans or two palpations to encourage them to adopt a constant arm elevation. The participants were not told that the reference stimulus was used in all trials and they did not receive any feedback. The standard stimulus was presented in every trial, and its position varied randomly on a trial-by-trial basis. The participant’s task was to indicated which of two consecutive shapes felt more curved, more stiff, or bigger. The curvature, stiffness and size of the comparison stimulus varied according to a one-up, one-down, interleaved staircase procedure which terminated after twenty reversals. The data for each condition and each participant were fit with cumulative Gaussian functions to extract PSEs and discrimination thresholds as exemplified in Fig. 6c with the results of participants S01 and S09.

## Data Availability

The data that support the findings of this study are available from the corresponding author upon reasonable request.

## Acknowledgments

The study was funded by the European Research Council (FP7) European Research Council Advanced Grant (Computational Theory of Haptic Perception) 247300 (to V.H.).

## References

[1] Koenderink, J. J. Solid shape (MIT Press, Cambridge, MA, 1990).

[2] Pont, S. C., Kappers, A. M. L. & Koenderink, J. J. Similar mechanisms underlie curvature comparison by static and dynamic touch. Perception, & Psychophysics 61, 874–894 (1999).

[3] Dostmohamed, H. & Hayward, V. Trajectory of contact region on the fingerpad gives the illusion of haptic shape. Experimental Brain Research 164, 387–394 (2005).

[4] van der Horst, J., B. & Kappers, A. M. L. Curvature discrimination in various finger conditions. Experimental Brain Research 3, 304–311 (2007).

[5] Sanders, A. F. J. & Kappers, A. M. L. Bimanual curvature discrimination of hand-sized surfaces placed at different positions. Perception, & Psychophysics 68, 1094–1106 (2006).

[6] Squeri, V. et al. Two hands, one perception: how bimanual haptic information is combined by the brain. Journal of Neurophysiology 107, 544–550 (2012).

[7] Cadieux, M. L., Whitworth, K. & Shore, D. I. Rubber hands do not cross the midline. Neuroscience Letters 504, 191–194 (2011).

[8] Bowers, D. & Heilman, K. M. Pseudoneglect: Effects of hemispace on a tactile line bisection task. Neuropsychologia 18, 491–498 (1980).

[9] Goldreich, D. A bayesian perceptual model replicates the cutaneous rabbit and other tactile spatiotemporal illusions. PLoS ONE 2, e333 (2007).

[10] Gross, G. C. & Mishkin, M. The neural basis of stimulus equivalence across retinal translation. Lateralization in the Nervous System 109–122 (1977).

[11] Funnell, M. G., Corballis, P. M. & Gazzaniga, M. S. Insights into the functional specificity of the human corpus callosum. Brain 123, 920–926 (2000).

[12] Erberich, S. G. et al. Somatosensory lateralization in the newborn brain. NeuroImage 29, 155–161 (2006).

[13] Barlow, H. Possible principles underlying the transformation of sensory messages. In Sensory Communication, 217–34 (MIT Press, Cambridge, MA, 1961).

[14] Holway, A. H. & Boring, E. G. Determinants of apparent visual size with distance variant. American Journal of Psychology 54, 21–37 (1941).

[15] Glennerster, A., Tcheang, L., Gilson, S. J., Fitzgibbon, A. W. & Parker, A. J. Humans ignore motion and stereo cues in favor of a fictional stable world. Current Biology 16, 428–432 (2006).

[16] Kong, G. et al. Interhemispheric communication during haptic self-perception. Proceedings of the Royal Society B 289, 20221977 (2022).

[17] Cattaneo, Z. et al. Spatial biases in peripersonal space in sighted and blind individuals revealed by a haptic line bisection paradigm. Journal of Experimental Psychology: Human Perception and Performance 37, 1110–1121 (2011).

[18] Yamamoto, S. & Kitazawa, S. Reversal of subjective temporal order due to arm crossing. Nature Neuroscience 4, 759–765 (2001).

[19] Yamamoto, S. & Kitazawa, S. Sensation at the tips of invisible tools. Nature Neuroscience 4, 979–980 (2001).

[20] Làdavas, E., Di Pellegrino, G. & Farné, G., A. amd Zeloni. Neuropsychological evidence of an integrated visuotactile representation of peripersonal space in humans. The Journal of Cognitive Neuroscience 10, 581–589 (1998).

[21] Fagot, J., Lacreuse, A. & Vauclair, J. Hand-movement profiles in a tactual—tactual matching task: Effects of spatial factors and laterality. Perception & Psychophysics 56, 347–355 (1994).

[22] Sutherland, M. T. & Tang, A. C. Reliable detection of bilateral activation in human primary somatosensory cortex by unilateral median nerve stimulation. Neuroimage 33, 1042–1054 (2006).

[23] Hagen, M. C. & Pardo, J. V. Pet studies of somatosensory processing of light touch. Behavioural Brain Research 135, 133–140 (2002).

[24] van Boven, R. W., Ingeholm, J. E., Beauchamp, M. S., Bikle, P. C. & Ungerleider, L. G. Tactile form and location processing in the human brain. Proceedings of the National Academy of Sciences 102, 12601–12605 (2005).

[25] Wijntjes, M., Sato, A., Hayward, V. & Kappers, A. M. L. Local surface orientation dominates haptic curvature discrimination. IEEE Transactions on Haptics 2, 94–102 (2009).

